# First records of invasive *Vespa velutina nigrithorax* Buysson, 1905 (Hymenoptera: Vespidae) and *Megachile sculpturalis* Smith, 1853 (Hymenoptera: Megachilidae) in Slovakia

**DOI:** 10.1101/2024.10.21.619466

**Authors:** Adrián Purkart, Marek Semelbauer, Peter Šima, Jozef Lukáš, Simon Hoffner, Peter Fedor, Dušan Senko

## Abstract

Biological invasions are an increasing threat to ecosystems; early identification of invasive species and rigorous monitoring are prerequisites to minimize environmental damage. Currently, two large hymenopterans of Asian origin are spreading across Europe: the yellow-legged hornet *Vespa velutina nigrithorax* Buysson, 1905 and the giant resin bee *Megachile sculpturalis* Smith, 1853, populations of which have been gradually being discovered across Europe since 2004 and 2008, respectively. Considering the current distribution of both species in Europe, further spread through Central Europe is expected in recent years. In July 2024, the first record of *M. sculpturalis* was documented in Slovakia, followed by more reports from 11 localities. Less than two months later, the second invasive hymenopteran, *V. velutina nigrithorax*, was also detected. Utilising multiple methods, their nest was discovered as well. On-site observations showed that the yellow-legged hornets (workers) were active almost two days after colony eradication. The finding of both species was accompanied by an intensive campaign using citizen science.

## Introduction

European nature is facing the intrusion of various non-native species of hymenopterans, which, by their way of life, threaten the native biota and can, therefore, be considered invasive. Such species include, e.g., the yellow-legged mud-dauber wasp *Sceliphron caementarium* (Drury, 1773), elm zigzag sawfly *Aproceros leucopoda* Takeuchi, 1939 or the Argentine ant *Linepithema humile* (Mayr, 1868) (Blank et al. 2010, López-Collar & Cabrero-Sañudo 2021, Bogusch 2022). One of the most iconic species that has successfully spread across Europe in the last 20 years is the yellow-legged hornet, *Vespa velutina nigrithorax* Buysson, 1905, which originates from Southeast Asia and was accidentally introduced to France in 2004 (Haxaire et al. 2006). This hornet is significantly smaller than the European hornet, *Vespa crabro* Linnaeus, 1758, while workers reach a size of approximately 20 mm. Another notable difference is that they are dark, with a typically yellow pattern on their abdomen and lighter yellow to orange legs. However, the biggest concern is their bionomics associated with the specialisation in hunting wide range of flying insects, including honey bees (*Apis mellifera* Linnaeus, 1758). If the *V. velutina nigrithorax* individuals discover a honeybee colony, they remember it as an easily accessible food source until they completely decimate it by gradual hunting the worker bees near their hives. By the end of summer 2023, this species was reported from Spain (Castro & Pagola-Carte 2010), Portugal (Grosso-Silva & Maia 2012), Belgium (Bruneau 2011), Italy (Demichelis et al. 2014), Germany (Witt 2015), Great Britain (Budge et al. 2017), the Balearic Islands (Leza et al. 2018), the Channel Islands (States of Guernsey Government 2016), the Netherlands (Smit et al. 2017), Switzerland (Ebener 2017), Luxemburg (Ries et al. 2021), Ireland (Dillane et al. 2022) and Hungary (Márta & Vas 2023). The observation in the Hungarian village of Kimle in the Győr-Moson-Sopron district prompted several communities to present various educational content on social networks and in the Czech, Slovak, and Austrian media (newspapers, TV, radio), aiming to use citizen science to discover them in the new nearby territories. As a result, new records of *V. velutina nigrithorax* were made in Austria (Schorkopf et al. 2024) and the Czech Republic (Walter et al. 2024).

Citizen science methods can also be applied to other noticeable species of the order Hymenoptera. One of them is the giant resin bee, *Megachile sculpturalis* Smith, 1853, a typical member of the family Megachilidae, whose native range spreads in the eastern Palaearctic (Roques et al. 2009). It belongs to the subgenus *Callomegachile* Michener, 1962, and is characterised by a large body (18–22 mm) and sizable mandibles adapted to excavate nest cavities in wood or reed stems. Over the past 25 years, *M. sculpturalis* has invaded new territories in North America (Mangum & Brook 1997) and Europe, where it was first time detected in France in 2008 (Vereecken & Barbier 2009), followed by Switzerland (Amiet, 2012), Italy (Quaranta et al., 2014), Germany (Westrich et al. 2015), Hungary (Kovács, 2015), Serbia (Ćetković & Plećaš 2017), Slovenia (Gogala & Zadravec, 2018), Austria and Liechtenstein (Dillier 2016; Westrich 2017; Lanner et al., 2020), Spain (Aguado et al., 2018), Ukraine (Ivanov & Fateryga, 2019), Bosnia and Herzegovina (Bila Dubaić et al., 2021) and the Mediterranean area (Ruzzier et al., 2020; Marquès & Calafat, 2021).

Based on previous findings, the further spread of these species in Europe was expected. In 2024, the great efforts of entomologists, in cooperation with the general public, resulted in the discovery of both invasive hymenopterans on the territory of Slovakia. This study presents the first faunistic records of their distribution across this region, supplemented with ecological notes.

## Material and methods

Anticipating the spread of the yellow-legged hornet to Slovakia, educational materials on its morphology and biology were translated into Slovak language. A website was also established in order to allow citizens to report suspicious insect findings (Diaz et al. 2022). These public awareness and environmental education activities led to the creation of additional educational materials in the form of leaflets, informational brochures, and local media. Each photograph received was reviewed and determined by entomologists within 24 hours. Following the first successful finds of both species, an intensive media and social media campaign was launched to gather additional faunistic data. Where possible, individuals were collected and deposited in the authors’ private collections and other relevant institutions.

Each locality of observation or collection contain the following information: Country name, geomorphologic unit, name of the locality (e.g., village, town, local site), faunistic code of the Central European mapping grid system (Ehrendorfer & Hamann 1965), locality identification (ID), coordinates (latitude, longitude), altitude, date of observation or collection, name of the insect, number of the observed/collected insects, sex or caste, name of the observer (obs.), collector (leg.), determinator (det.), observation details such as trophic interactions, specific remarks and collection methods.

Photographs of voucher specimens were taken on a Leica M205 C and edited with Adobe Photoshop. The data map was created in ArcMap 10.8 (Esri 2020) and included a combination of the data presented in this study and data obtained from the iNaturalist portal (iNaturalist 2024a,b), and GBIF (GBIF.org 2024). The botanical nomenclature follows SlovPlantList – the database of the names of vascular plants of Slovakia (Letz et al. 2024).

## Results

### *Vespa velutina nigrithorax* Buysson, 1905

On September 28, 2024, several yellow-legged hornets were discovered in the village of Palárikovo (Nové Zámky district) by Plant Science and Biodiversity Centre Slovak Academy of Sciences (PSBC) employees. On September 30, 2024, the State Nature Conservancy of the Slovak Republic, in collaboration with PSBC, conducted various methods, including triangulation (see Leza et al. 2018), to locate the nest. On October 1, 2024, Dr. Balázs Kolics (Hungarian University of Agriculture and Life Sciences) and Dr. Helena Proková (Blesabee) joined the team, using VHF radiotelemetry to track the hornets’ movements. Transmitters were glued to the hornets, transmitting a VHF signal at a specific frequency. Tracking was conducted by following the signal strength and direction until the nest’s proximity was indicated.

Slovakia, Podunajská rovina, Palárikovo, 7974, 48°02’07.0”N 18°04’14.6”E (Fig. 1,2,3), locality ID P1, P2, 110 m a. s. l., 28.09.2024, *Vespa velutina nigrithorax* (Fig. 4), up to 10 females (workers), leg. det. et coll. K. Senková Baldaufová & D. Senko, observed on fallen apples and 32 m further on the five inflorescences of the banana plant (*Musa basjoo*) in the private garden, collected by an entomological net. The very first observation of the species in Slovakia.

**Fig. 1:**
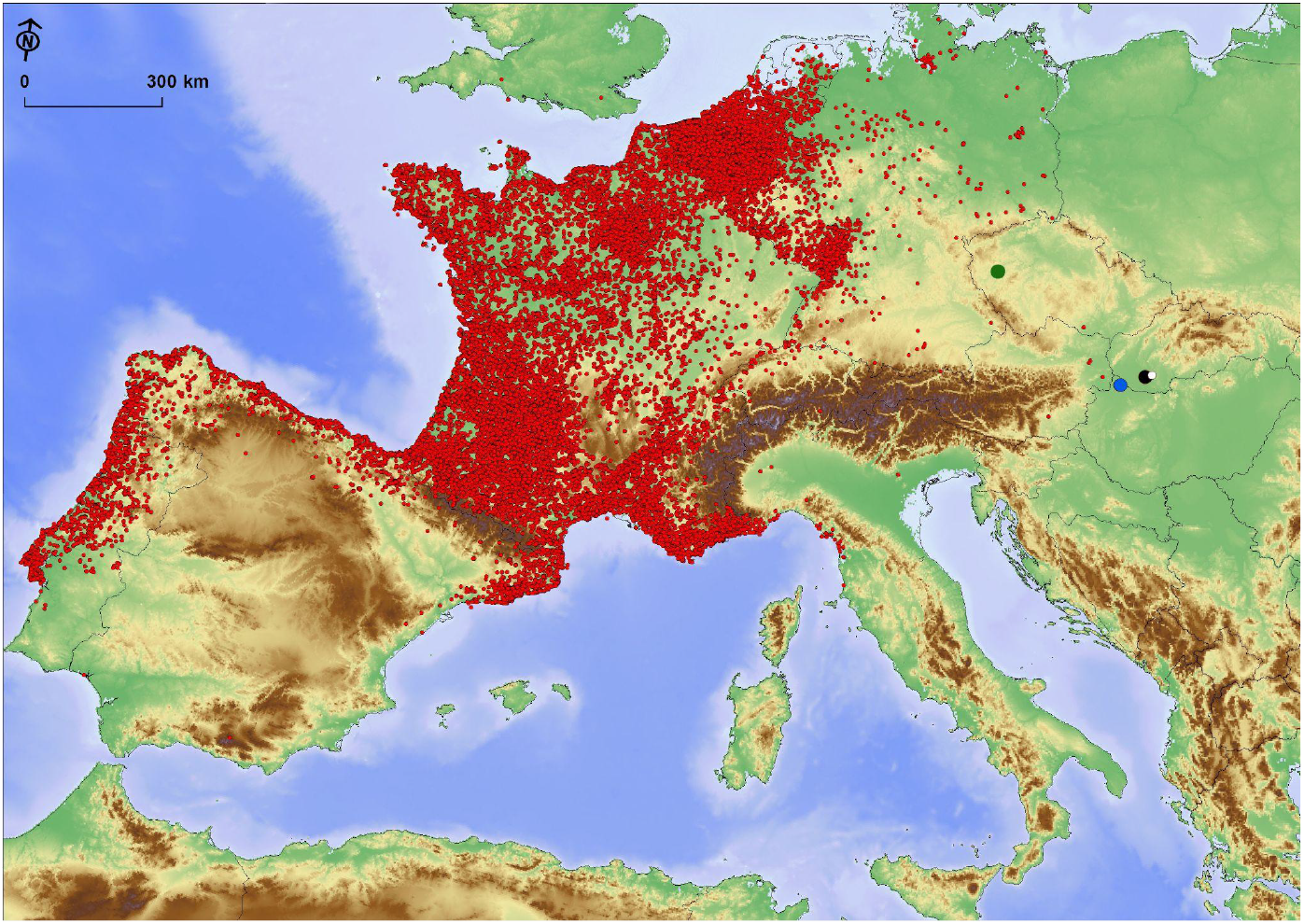
Slovak observations of *Vespa velutina* within Europe. Showing data obtained from the iNaturalist portal and GBIF. The larger dots on the map represent confirmed nests in Czech (green), Hungary (blue), and Palárikovo (black). White dot is Dolný Ohaj.

**Fig. 2:**
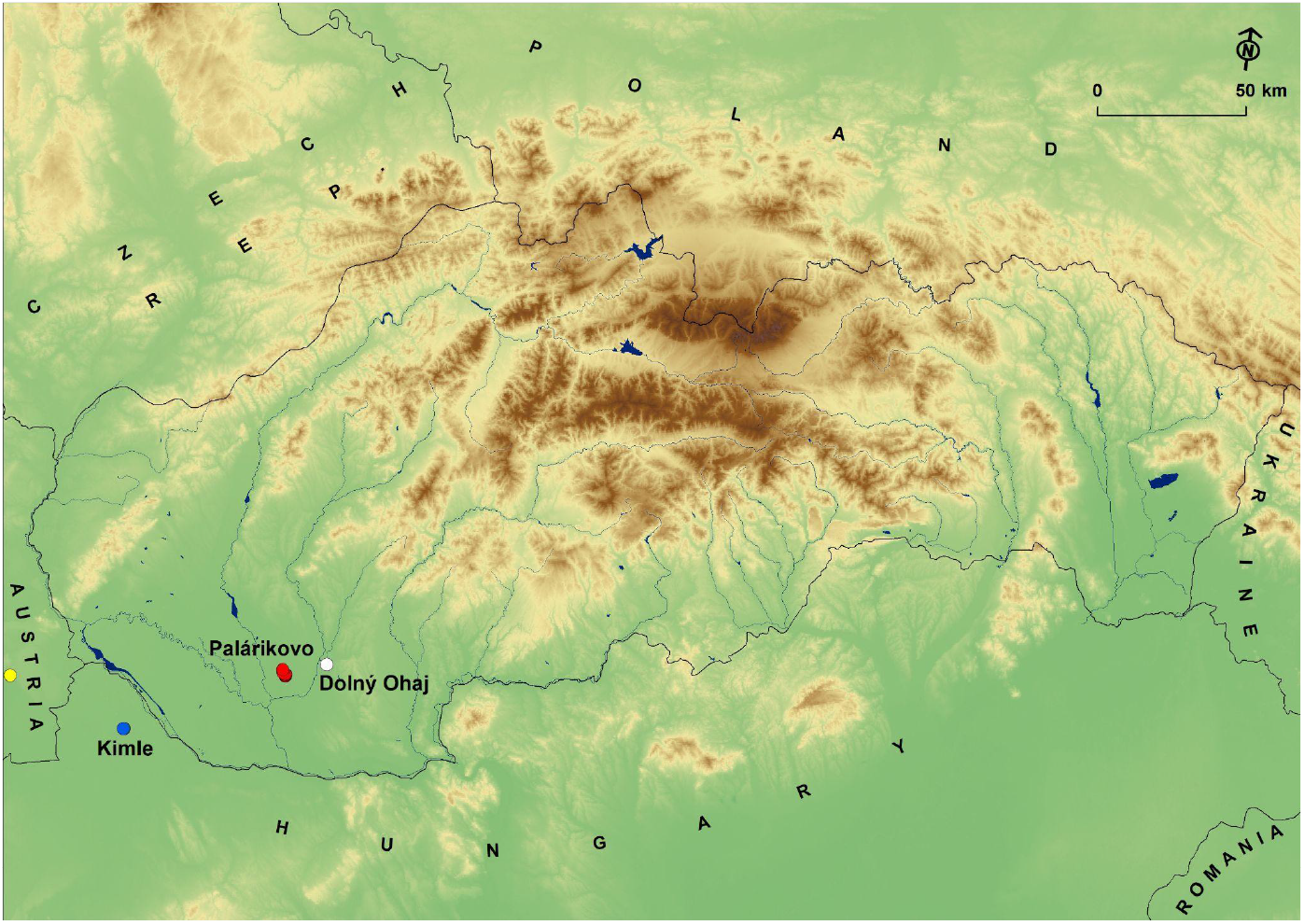
Observations of *V. velutina* in Palárikovo and Dolný Ohaj. Kimle (blue) in Hungary is also shown (the distance to Palárikovo (red) is 57.2 km). Dolný Ohaj (white): The difference between Slovak observations is 14.5 km. The yellow dot represents an observation in Austria from May 21, 2023.

**Fig. 3:**
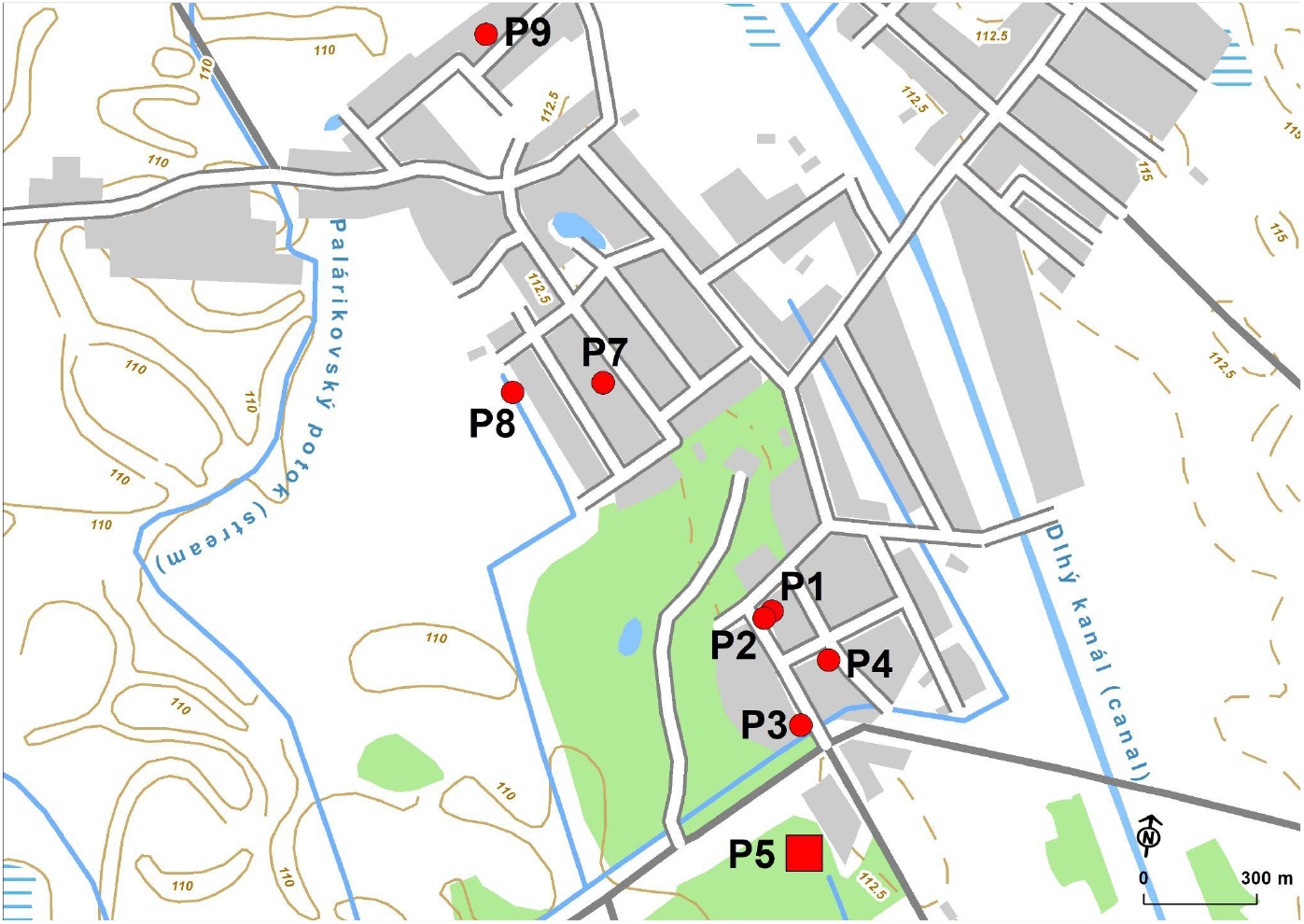
Detailed localization of occurrences of *V. velutina* in Palárikovo. P1: first observation on fallen apples, P2: *Musa basjoo*, P3: *Hedera helix* on J. Kalinčiaka st., P4: *Hedera helix* on Čs. armády st., P5: nest, P6: Hlavná st. (observer J. Húska), P7: Janošíková st. (observer F. Cirók), P8: Mederská st. (observer R. Vitko)

**Fig. 4:**
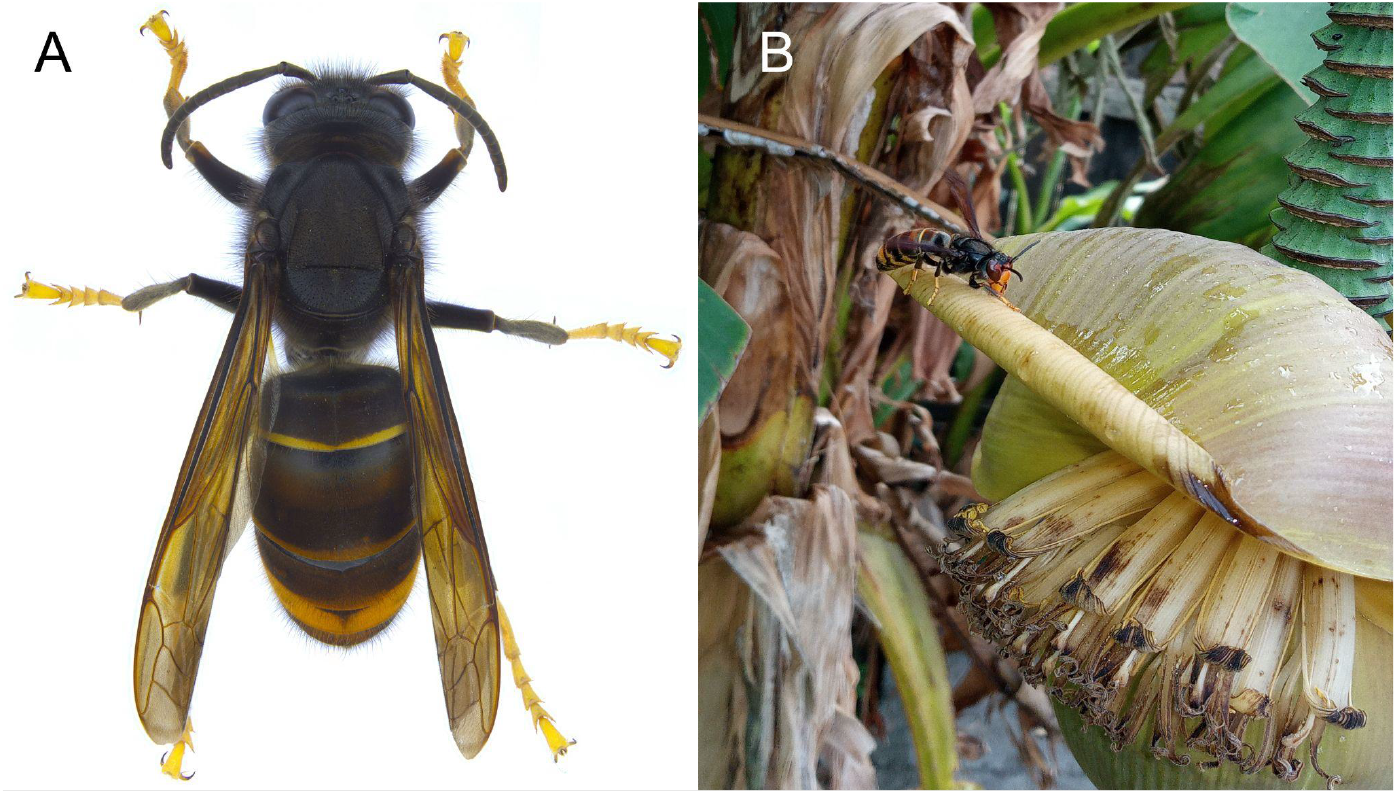
A - *Vespa velutina nigrithorax* habitus, worker, dorsal view; B - *V. velutina nigrithorax* worker visiting flower of cultivated banana, Palárikovo.

Slovakia, Podunajská rovina, Palárikovo, 7974, 48°01’57.3”N 18°04’21.6”E (Fig. 3), locality ID P3, 110 m a. s. l., 30.09.2024, *Vespa velutina nigrithorax*, up to 20 females (workers), leg. det. et coll. D. Senko, on inflorescences of *Hedera helix* in the private garden, collected by an entomological net.

Slovakia (SW), Podunajská rovina, Palárikovo, 7974, 48°02’03.1”N 18°04’24.3”E (Fig. 3), locality ID P4, 111 m a. s. l., 01.10.2024, *Vespa velutina nigrithorax*, up to 5 females (workers), leg. det. et coll. D. Senko, on inflorescences of *Hedera helix* in the private garden, collected by an entomological net.

On October 3, 2024, at 17:59, employees of pest control company Aquazoo s.r.o., using a drone with an infrared camera (see Walter et al. 2024), detected in an area that localized the methods of trigonometry and telemetry a temperature anomaly (+2.5 to 3°C compared to the 13°C ambient air temperature). This led to the discovering of the first-ever *Vespa velutina nigrithorax* nest in Slovakia, which Dr. Jozef Lengyel spotted. The nest was located on a grey poplar (*Populus × euroamericana*) at a height of approximately 22 m, and its size was around 80 cm in diameter (48°01’46.3”N 18°04’23.5”E, Fig 3,5), locality ID P5). It was found in the largest pheasantry in Central Europe, in the local part of the village called India. The species composition of the nest’s surrounding area can be classified as a hard riparian forest (Oak-elm and ash forests), characterized by waterlogged soils due to a high groundwater level. It is an open and sparse stand, lacking a developed shrub layer. In the herbaceous layer, *Rubus caesius, Carex hirta*, and *Geranium robertianum* are dominant. The tree species primarily include *Quercus robur* agg. and *Fraxinus excelsior*. The stand itself is estimated to be about 25 years old, while the tree with the nest, *Populus × euroamericana*, is estimated to be 25-30 years old, with a circumference at breast height of 135 cm. The tree is located at the edge of the stand. The nest was situated 663 m from the first recorded observation of specimens. The hornets were eradicated using insecticides on October 7, 2024, at 18:15. The tip pierced 40 cm into the lower part of the nest (Fig. 5). A field survey on 9 and 11 October in the vicinity of the nest of *V. velutina nigrithorax* revealed a suspicious large hornet nest in a tree near the village of čechy (48°02’29.7”N 18°22’31.7”E). Closer observation showed it was a *Vespa crabro* nest (Fig. 5).

**Fig. 5:**
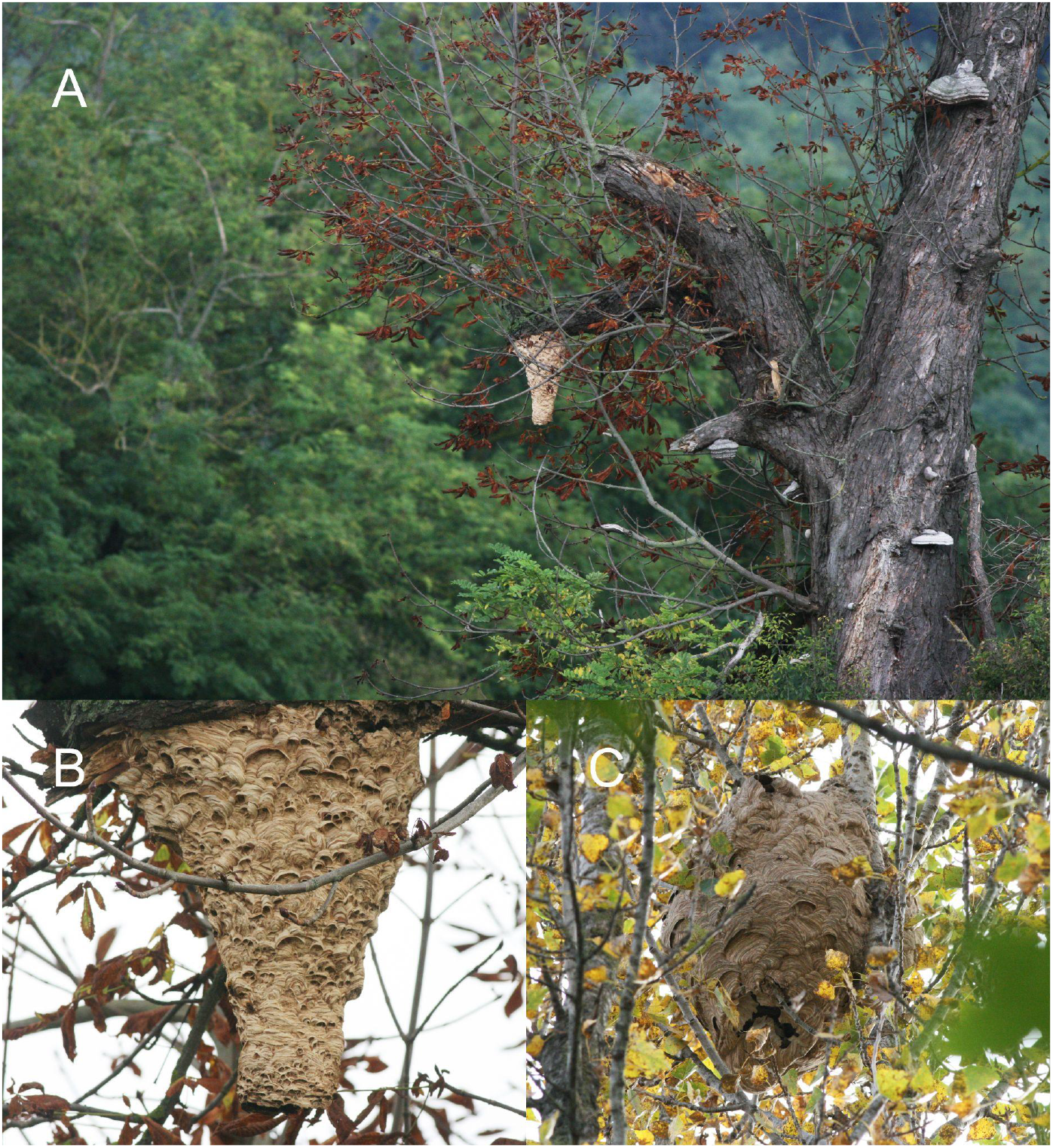
Hornet nests. A - *Vespa crabro* nest on European horse-chestnut tree, B - *Vespa crabro* nest, detail, C - *Vespa velutina nigrithorax* nest photographed from the ground, Palárikovo.

The entire 25-m tree with the nest was felled on October 17, 2024. The nest was broken into unknown pieces upon impact. On Monday, October 21, the nest was transferred from Banská Bystrica to the Faculty of Natural Sciences of Comenius University in Bratislava, where its fragments were deposited for further research.

Slovakia, Podunajská rovina, Palárikovo, 7974, 48°1’57.3”N 18°4’21.6”E (Fig. 3), locality ID P3, 110 m a. s. l., 05.10.2024, *Vespa velutina nigrithorax*, 20 females (workers), leg. det. et coll. P. Šima & A. Purkart, on inflorescences of *Hedera helix* in the private garden; 06.10.2024, same locality 1 female (worker), leg. det. et coll. P. Šima, on inflorescences of *Hedera helix* in the private garden, 341 m from the nest, collected by an entomological net.

Slovakia, Podunajská rovina, Palárikovo, 7974, 48°02’25.1”N 18°03’52.4”E (Fig. 3), locality ID P7, 110 m a. s. l., 06.10.2024, *Vespa velutina nigrithorax* (Fig. 6), 2 females (workers), leg. det. et coll. J. Húska, on sliced apples in the private garden, 1359.5 m from the nest, specimen collected by a jar.

**Fig. 6:**
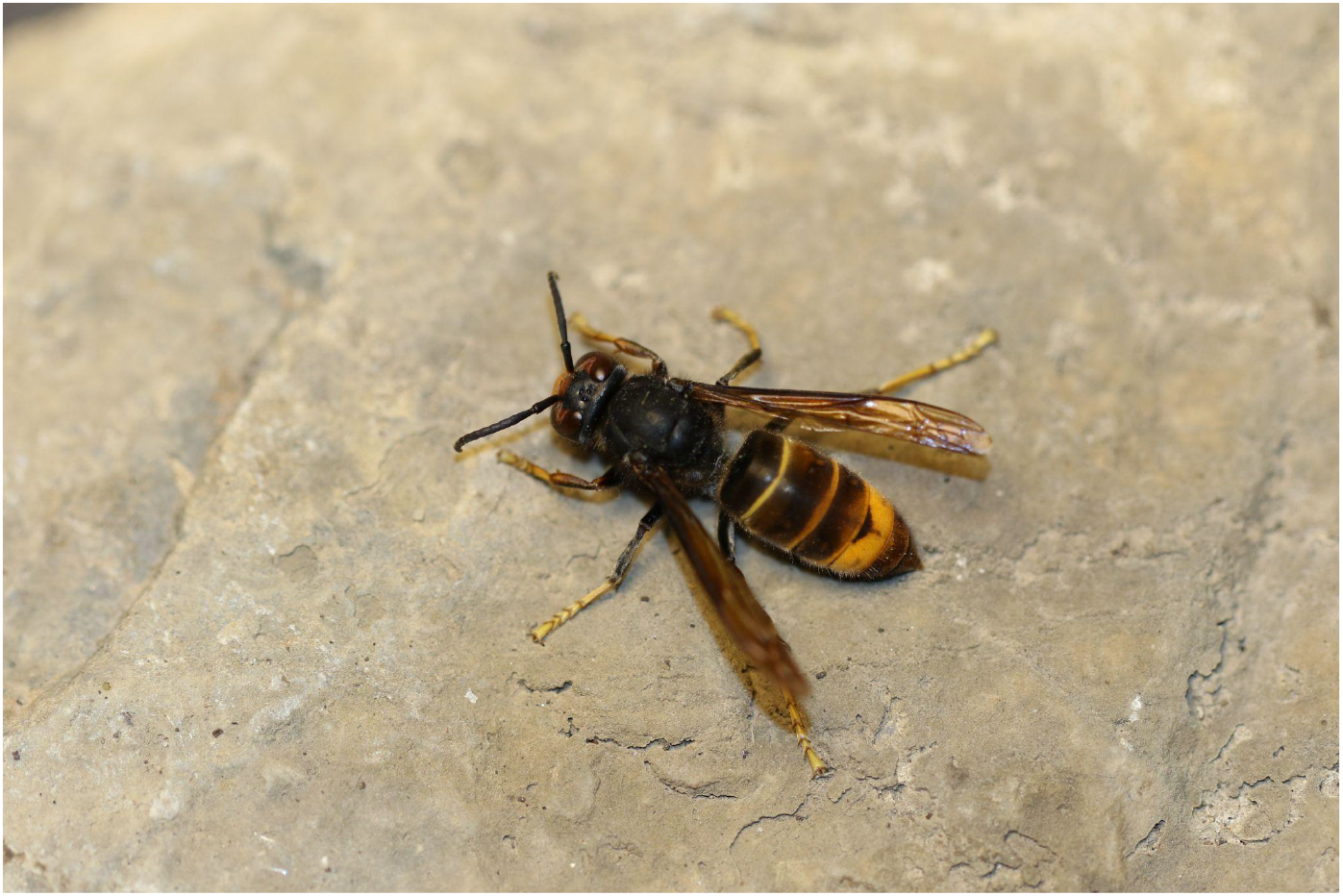
*Vespa velutina nigrithorax* worker from Palárikovo.

Slovakia, Podunajská rovina, Palárikovo, 7974, 48°02’23.7”N 18°03’40.9”E (Fig. 3), locality ID P8, 108 m a. s. l., 08.10.2024, *Vespa velutina nigrithorax*, 1 male, leg. det. K. Senková Baldaufová, coll. F. Cirók on the wall of a family house, 1.452 m NNW from the nest (cca 21 hours after nest eradication), specimen collected by a jar.

Slovakia, Podunajská rovina, Palárikovo, 7974, 48°01’57.3”N, 18°04’21.6”E (Fig. 3), locality ID P3, 110 m a. s. l., 08.10.2024, *Vespa velutina nigrithorax*, 5 females (workers), leg. det. M. Senko, flying around *Hedera helix*, one specimen was observed hunting a honeybee (*Apis mellifera*), 341 m from the nest (cca 22 hours after the extermination process).

Slovakia, Podunajská rovina, Palárikovo, 7974, 48°01’46.3”N 18°04’23.5”E (Fig. 3), locality ID P5, 110 m a. s. l., 09.10.2024, *Vespa velutina nigrithorax*, a few females (workers) and up to 10 males, leg. det. et coll D. Senko, observed beneath the nest, moving slowly and gradually dying; around the nest, there was no movement; no individuals were observed at several known localities within the urban area (cca 46 hours after the extermination process)

Slovakia, Podunajská rovina, Palárikovo, 7974, 48°02’54.0”N 18°03’34.0”E (Fig. 3), locality ID P11, 113 m a. s. l., 09.10.2024, *Vespa velutina nigrithorax*, 1 female (worker), leg. det. K. Senková Baldaufová, coll R. Vitko, on the bunches of grapes (*Vitis vinifera*) in the private garden, 2.328 m NNW from the nest (47,5 hours after the extermination process), this specimen was the last known observation in the vicinity of the nest site

Slovakia, Podunajská rovina, Palárikovo, 7974, 48°01’46.3”N 18°04’23.5”E (Fig. 3), locality ID P5, 110 m a. s. l., 11.10.2024, *Vespa velutina nigrithorax*, 14 females (workers) and 20 males, leg. det. et coll. D. Senko, dead specimens collected around the treated nest

Slovakia, Hronská pahorkatina, Dolný Ohaj, 7975, 48°04’33.2”N 18°15’20.9”E (Fig. 2), 122 m a. s. l., 15.10.2024, 14:30, *Vespa velutina nigrithorax*, 1 male, leg. R. Gatyáš, det. P. Šima, coll. D. Senko, feeding on a pear fruit, 14.5 km from the nest in Palárikovo, specimen collected by a jar.

### *Megachile sculpturalis* Smith, 1853

Slovakia, Borská nížina, Bratislava, 7767, 48°12’24.3”N 16°58’30.2”E (Fig. 7), 155 m a. s. l., 08.7.2024, *Megachile sculpturalis*, 1 female, leg. det. et coll. J. Lukáš, collected on *Sophora japonica*.

**Fig. 7:**
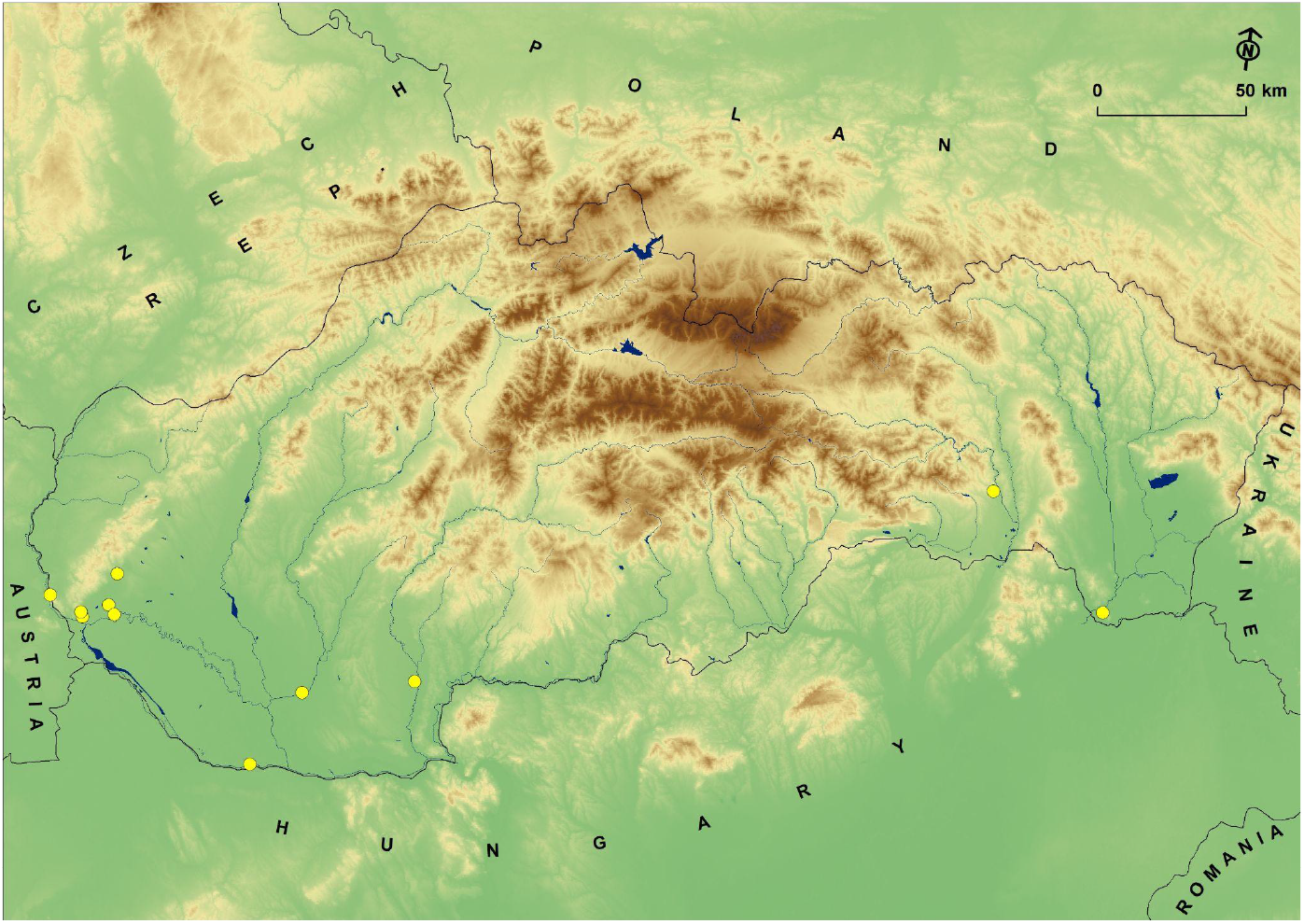
Observations of *Megachile sculpturalis* in Slovakia.

Slovakia, Východoslovenská rovina, Streda nad Bodrogom, 7696, 48°22’52.5”N 21°45’47.7”E (Fig. 7), 125 m a. s. l., 9.7.2024, 1 female, leg. Štefan Timko, det. M. Semelbauer, specimen observed in an apiary.

Slovakia, Podunajská rovina, Bratislava, 7868, 48°09’02.9”N 17°07’52.6”E (Fig. 7), 144 m a. s. l., 12.7.2024, *Megachile sculpturalis*, 1 female, leg. det. et coll. J. Lukáš, found on a concrete wall close to *Sophora japonica*.

Slovakia, Košická kotlina, Košice, 7293, 48°44’07.4”N 21°14’18.7”E (Fig. 7), 223 m a. s. l., 18.7.2024, *Megachile sculpturalis*, 1 female, leg. S. Semešová, det. A. Purkart, observed in a botany garden

Slovakia, Podunajská rovina, Ivanka pri Dunaji, 7869, 48°11’40.8”N 17°14’38.6”E (Fig. 7), 155 m a. s. l., 21.7.2024, *Megachile sculpturalis*, 1 female, leg. det. et coll. J. Lukáš, collected on *Sophora japonica*

Slovakia, Podunajská rovina, Veľký Lel, 8273, 47°45’11.7”N 17°56’52.8”E (Fig. 7), 115 m a. s. l., 26.7.2024, *Megachile sculpturalis*, 1 female (Fig. 2), leg. S. Hoffner det. et coll. M. Semelbauer, captured in a Danube floodplain habitat, Fig. 8.

**Fig. 8:**
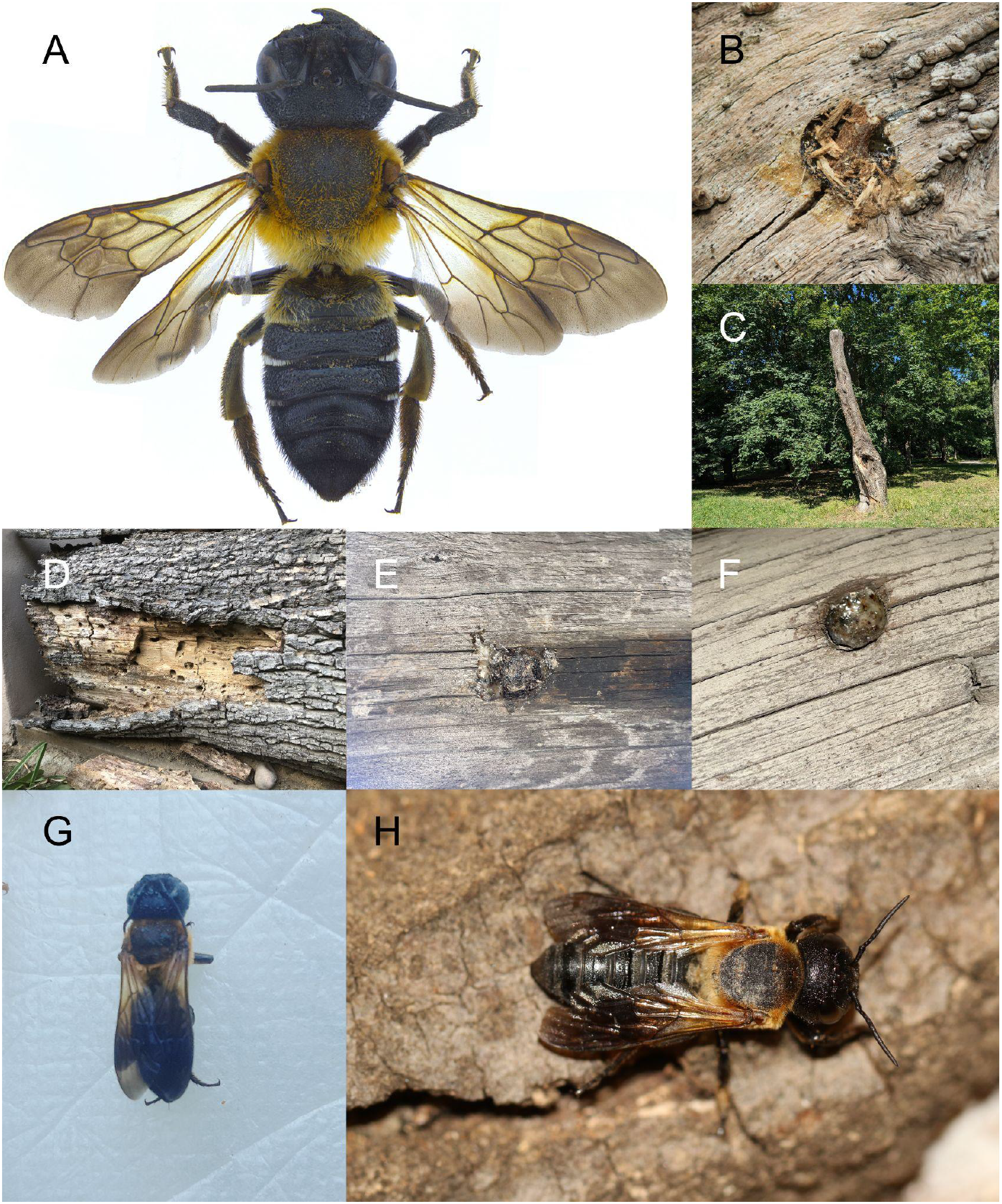
Habitus, nests and habitats of *Megachile sculpturalis* A - *Megachile sculpturalis* habitus, female, dorsal view; B - nest of *M. sculpturalis*, Pezinok city park; C - Dead tree trunk with nests of *M. sculpturalis* in city par, Pezinok; D - Old walnut tree (*Juglans regia*) trunk with a large nesting population of *M. sculpturalis* (Želiezovce); E, F - Nesting holes of *M. sculpturalis* filled with resin (Nové Zámky); G - *M. sculpturalis* female observed in an apiary, Streda nad Bodrogom; H - *M. sculpturalis* female, Želiezovce.

Slovakia, Podunajská rovina, Zálesie, 7869, 48°10’00.5”N 17°16’25.7”E (Fig. 7), 130 m a. s. l., 26.7.2024, *Megachile sculpturalis*, 1 female, leg. det. et coll. A. Purkart, captured in a garden of a family house

Slovakia, Hronská pahorkatina, Želiezovce, 7977, 48°2’47”N 18°39’30”E (Fig. 7), 134 m a. s. l., 07. - 08.2020, *Megachile sculpturalis*, >10 females, observ. J. Sýkorčin, females were detected during nesting in an old trunk of *Juglans regia* in a private garden. The landowner mentioned that he has been observing this species there since at least 2020.

Slovakia, Hronská pahorkatina, Želiezovce, 7977, 48°2’47”N 18°39’30”E (Fig. 7), 134 m a. s. l., 10.08.2024, *Megachile sculpturalis*, 5 females, leg. det. et coll. P. Šima, 2 females leg. et det. P. Šima, coll. V. Smetana, captured in a private garden, nesting in an old trunk of *Juglans regia* (Fig. 8).

Slovakia, Podunajská rovina, Nové Zámky 8074, 47°59’03”N 18°9’17”E (Fig. 7), 113 m a. s. l., 15.08.2024, *Megachile sculpturalis*, 1 female, leg. J. Urbán, det. et coll. P. Šima, private garden, nesting in an old wooden barn construction (Fig. 8).

Slovakia, Podunajská rovina, Pezinok, 7769, 48°17’29.9”N 17°16’07.5”E (Fig. 7), 164 m a. s. l., 02.8.2024, *Megachile sculpturalis*, 1 female, leg. V. Marušic, det. A. Purkart, observed building a nest in an old tree trunk in the castle park (Fig. 8).

Slovakia, Podunajská rovina, Bratislava, 7869, 48°09’55.3”N 17°07’22.2”E (Fig. 7), 150 m a. s. l., 19.8.2024, *Megachile sculpturalis*, 1 female, leg. S. Felcan, det. A. Purkart, observed on a wooden rooftop of a reconstructed building.

## Discussion

The yellow-legged hornet (*Vespa velutina*) originates from Southeast Asia and was accidentally introduced to France in 2004 (Haxaire et al. 2006), spreading rapidly across Europe. In Hungary, they were confirmed on August 19, 2023, in the village of Kimle in the Győr-Moson-Sopron district, 57.2 km from the first finding in Palárikovo, Slovakia. Two workers were found in a local apiary (Márta & Vas 2023). Later, the nest was tracked using radiotelemetry and destroyed after 48 days (Vas, pers. comm.). In the Czech Republic, the yellow-legged hornets were detected in Pilsen on October 5, 2023 (Walter et al. 2024) on the black locust tree (*Robinia pseudoacacia*). The nest was discovered using a triangulation method (Leza et al. 2018) on October 9, 2023, and subsequently removed. It was 395.39 km from the Slovakian nest. In Austria, the species was discovered in Salzburg on April 9, 2024 (Schorkopf et al. 2024).

The search for the first Slovak nest of *V. velutina* was carried out by the relevant government authorities in cooperation with several experts. Similar to the experience from other European countries, the method of telemetry and triangulation proved successful in pinpointing the area where the nest may be located. Franklin et al. (2017) describe that over 90 % of *V. velutina* nests are located in the tree canopy, which was also confirmed in this case. The nest was located on the edge of a forest stand and close to water sources: 197 m from the Žofiin canal and 286 m from a local pond. Similar observations were made in the Czech Republic, where the nest was found 80 metres from the Vejprnický stream. This confirms the results of Monceau et al. (2012) that *V. velutina* prefers to build nests in landscapes with abundant water resources. The first nest in the Czech Republic has also been thoroughly analysed (Walter et al. 2024), providing valuable insights into the bionomics of this invasive species in this part of Europe where it is expanding. In the case of the nest in this study, it was not technically simple to gently remove it from the tree canopy and was only obtained 10 days after insecticide application, which resulted in its condition being insufficient for further laboratory analyses. In contrast, valuable observations were obtained in the case of the workers. These were found only in the village of Palárikovo, where they searched for food with high sugar content. On inflorescence of *H. helix*, they primarily fed on nectar together with other insect species (honey bees, wasps, and hoverflies), which yellow-legged hornets also actively hunted. Species identification of the prey, except for the honey bee in one case, was not determined because the hunting took place out of visual range. The yellow-legged hornet is a well-known threat to beekeeping (Perrard et al. 2009), but its impacts on biodiversity and wild pollinators are much less studied. Workers were still active near the site approximately two days after its eradication by insecticide. One male was also found 14.53 km from the nest eight days after this process. After treatment, a closer examination of the dead individuals confirmed that the males had already hatched in the nest and may have been active in the area surrounding the nest site. Young hatched queens of *V. velutina nigrithorax* were not detected. The spread of this species in the Central European region is not surprising, and the ability to establish populations here is suggested by the studies of Abou-Shaara & Al-Khalaf (2022) and Walter et al. (2024). The dispersal rate here is yet unknown. The estimated invasion speed of *V. velutina* differs considerably among the countries of occurrence. The spread rate was estimated to be about 78 km/year in France (Robinet et al. 2017), while in Italy (Liguria region), it was 18,3±3,3 km/year (Bertolino et al. 2016), and in Portugal it varies from 20 - 45 km/year (Carvalho et al. 2020, Verdasca et al. 2021) The differences are most probably influenced by the climatic suitability of the region, initial population density, landscape topography and elements, phytocoenosis of the region and road infrastructure, among others. Nest densities, when considering both the urban and non-urban areas, may reach 2,9 - 4,81 nests/km^2^ (Bertolino et al. 2016, Franklin et al. 2017)

Close collaboration between citizen science and scientists has resulted in the early detection of invasive species. In the case of the yellow-legged hornet in Slovakia, continuous monitoring was carried out using data sent by the general public to the staff of The State Nature Conservancy of the Slovak Republic using the online portal and mobile application Zastav sršňa (“Stop the Hornet”). Photographs of various organisms that resembled *V. velutina nigrithorax* in appearance were submitted, including many findings of *Megascolia maculata* (Drury, 1773), which is a rare insect species in Slovakia. Other insect species with which the public confused the yellow-legged hornet were, for example, hoverflies of the genus *Volucella*, robber flies of the family Asilidae, or even hummingbird hawk-moth of the species *Macroglossum stellatarum* (Linnaeus, 1758).

Similar principles of data collection using citizen science have been applied to a second invasive species of hymenopteran, *Megachile sculpturalis*. In this case, as many as 6 of the 12 records were found thanks to the public, which only demonstrates the importance of this data-gathering method in the field. Its large size is also beneficial in monitoring the occurrence of this species. Dubaić and Lanner (2021) add that beekeepers are an especially appropriate group to map this species because, unlike most of the public, they can recognize the honey bee from the other solitary bees, including the giant resin bee. The presented findings also confirm the further expansion of *M. sculpturalis* in Central Europe. Given the uniform character of the Pannonian Basin and Danube Plain, continued spreading is to be expected. Lanner et al. (2020) found that in the Alpine region, these solitary bees can colonise habitats up to 1230 m altitude. Despite the mountainous nature of Slovakia, the distribution could thus cover almost the whole country in the future and potentially reach Poland and other northern countries. This species of giant resin bee is considered invasive because females often use already existing nests of different bee species (e.g., *Osmia* sp.) during the establishment of their nest cavities. However, it is not clear from observations to date whether this is a rare occurrence or a form of direct or indirect competition between *M. sculpturalis* and native cavity nesters (Roulston & Malfi 2012; Lanner et al. 2020). For this reason, it is also worth tracing its further spread across the region. The easiest way to look for this species is to observe ‘bee hotels’ and old tree trunks, where females like to occupy holes with a diameter of 8–20 mm (Le Féon et al. 2018). We consider it appropriate to encourage the general public in mapping as well, perhaps with the help of social media. Applications like iNaturalist.org or observation.org are also helpful, as people can contribute observations, and at the same time, their observations are determined by other users. Invasive species are often large and conspicuous, and the public usually reacts strongly to invasions (as a threat to beekeeping and biodiversity), providing an opportunity to draw the general public’s attention to wild pollinators.

## Acknowledgment

We are very grateful to Mgr. Katarína Senková Baldaufová (Plant Science and Biodiversity Centre, Slovak Academy of Sciences) for her initial discovery of *V. velutina nigrithorax* in Palárikovo. We would like to thank all citizen scientists, Jozef Lengyel, Richard Schnieder, J. Sýkorčin, J. Urbán, A. Urbánová, F. Cirók, J. Húska, R. Gatyáš, R. Vitko, Š. Timko, S. Semešová, S. Felcan, V. Marušic, and the State Nature Conservancy of the Slovak Republic employees for their help collecting samples in the field. We altho thank Ľubomír Vidlička (Institute of Zoology, Slovak Academy of Sciences) for preparing high-quality images of the two invasive species. The study was financially supported by the project VEGA 2/0022/23, 1/0372/24, and LIFE 14 NAT/SK/001306 Restoration and management of Danube floodplain habitats.

